# Quantitative Geometric Modeling of Blood Cells from X-ray Histotomograms of Whole Zebrafish Larvae

**DOI:** 10.1101/2023.05.23.541939

**Authors:** Maksim A. Yakovlev, Ke Liang, Carolyn R. Zaino, Daniel J. Vanselow, Andrew L. Sugarman, Alex Y. Lin, Patrick J. La Riviere, Yuxi Zheng, Justin D. Silverman, John C. Leichty, Sharon X. Huang, Keith C. Cheng

## Abstract

Tissue phenotyping is foundational to understanding and assessing the cellular aspects of disease in organismal context and an important adjunct to molecular studies in the dissection of gene function, chemical effects, and disease. As a first step toward computational tissue phenotyping, we explore the potential of cellular phenotyping from 3-Dimensional (3D), 0.74 µm isotropic voxel resolution, whole zebrafish larval images derived from X-ray histotomography, a form of micro-CT customized for histopathology. As proof of principle towards computational tissue phenotyping of cells, we created a semi-automated mechanism for the segmentation of blood cells in the vascular spaces of zebrafish larvae, followed by modeling and extraction of quantitative geometric parameters. Manually segmented cells were used to train a random forest classifier for blood cells, enabling the use of a generalized cellular segmentation algorithm for the accurate segmentation of blood cells. These models were used to create an automated data segmentation and analysis pipeline to guide the steps in a 3D workflow including blood cell region prediction, cell boundary extraction, and statistical characterization of 3D geometric and cytological features. We were able to distinguish blood cells at two stages in development (4- and 5-days-post-fertilization) and wild-type vs. *polA2 huli hutu* (*hht*) mutants. The application of geometric modeling across cell types to and across organisms and sample types may comprise a valuable foundation for computational phenotyping that is more open, informative, rapid, objective, and reproducible.

## 1. Introduction

Structurally distinct microscopic characteristics of eukaryotic cells allow the recognition of cell types and the study of physiological and pathological change. The recognition of cellular pathology [Virchow, 1860], while largely qualitative and subjective, has played a critical role in patient care for over 100 years (Abbas et al., 2015). Light microscopy, whose resolution at commonly used magnifications is in the 0.5 to 1 micron range, has been the most common modality used for distinguishing cells and their features. Histological analysis allows for the characterization of normal and abnormal tissue structure including cellular-resolution scale features and the study of the arrangement of cells into epithelial, stromal, and other structures.

Attempts to characterize these micron-scale structures in 3-Dimensional (3D) Euclidean space (Cheng et al., 2011) have included serial sectioning in a consistent plane of section (Hewitson & Darby, 2010). However, this approach is associated with sectioning artifact (Flaherty et al., 2001) (Arganda-Carreras et al., 2010). Ideally, 3D phenotyping would be done without physical sectioning to minimize sample distortion (Hillman, 2000) associated with cutting, tissue shrinkage associated with fixation, and differential expansion associated with floating sections on water (Kushida, 1962). In response to the growing need for 3D cellular-resolution imaging, modalities including electron microscopy, fluorescence imaging, magnetic resonance imaging (MRI), optical projection tomography, optoacoustic microscopy, and micro-computed tomography (micro-CT) have been applied towards closing the mesoscale imaging gap in model organisms such as zebrafish (Harris et al., 2006), (Frangioni, 2003), (Cramer & Rust, 1986), (Metscher, 2009), (Vinegoni et al., 2009) (Zhang et al., 2006), (Cheng et al., 2011). High resolution methods like serial-section electron microscopy (ssEM) can produce nanometer-scale slices that can be stacked into high-resolution 3D datasets, but are resource-heavy techniques that needs both a long time commitment as well as computer infrastructure capable of analyzing terabyte-scale data per scan (Hildebrand et al., 2017). In contrast, modalities like MRI or functional MRI are capable of scanning samples in minutes and generate significantly smaller datasets, but the lower resolution prevents cellular-level analysis (Dodd et al., 1999), (Flogel & Ahrens, 2016). To bridge this gap between resolution, field of view, and analytical feasibility, micro-CT, which can provide isotropic submicron-scale resolution (Mizutani et al., 2013), is necessary for millimeter to centimeter scale samples (Ding et al., 2019), (Yakovlev et al. 2022).

Histotomography, a form of micro-CT customized for histopathology (Ding et al., 2019), is based on tomographic reconstructions from series of angular X-ray projections of metal-stained tissue samples, providing about 1000-fold higher resolution than clinical CT scans. Functionally important geometric characteristics of cells and tissue structures such as volume and 3D shape, are distinguishable from such X-ray histotomographic images at submicron voxel resolutions. In addition to addressing the need for micron-scale imaging, micro-CT is a nondestructive imaging modality that allows rescanning of samples multiple times under different imaging conditions. Access to the entire tissue or animal sample ensures the preservation of all information inherently available in the specimen but presents the issue of characterizing cytological details in datasets harboring upwards of thousands of cells. Whole-sample 3D imaging using histotomography avoids incomplete sampling and begins to make possible phenotyping of whole organisms. In earlier work, we were able to show visual differences in the phenotype of a mutant vs. wild-type zebrafish (Ding et al. 2019). Here, we begin to explore the potential of modeling cells to quantify geometric shape parameters of individual cell types from whole-organism histotomograms.

The growing community demand to scan larger samples at finer levels of detail has facilitated the improvement of resolution and field of view in micro-CT imaging while simultaneously creating a need for better tools to interrogate large datasets. Image acquisition and data processing techniques such as phase-contrast imaging, mosaic spiral-CT acquisition, dual-energy scanning, and post-acquisition software-based processing tools such as distortion correction have allowed micro-CT imaging to overcome limitations of detector technologies through effective, but computationally expensive solutions (Barbone et al., 2021), (Silva et al., 2017), (Sawall et al., 2012), (Vo et al., 2015), (Pelt & Parkinson, 2018). Alternatively, the development of higher-resolution, wider field-of-view optics systems achieves this imaging goal directly and independently of any of the previously mentioned unconventional acquisition approaches. Multiple research groups, including our own, have adapted commercial high-mega-pixel cameras alongside custom-built objective lenses for micro-CT, enabling centimeter scale imaging with micron-scale detail at resolution to field-of-view ratios of up to 1:10,000 (Umetani et al., 2020), (Yakovlev et al., 2022). As these methodologies develop to generate larger and larger 3D image datasets, the need for accessible tools capable of analyzing the data beyond visualization of representative slices proportionately increases. Likewise, the increase in data volume makes manual processes, such as manual segmentation of volumes of interest, prohibitive and time consuming. Automated tools and data pipelines have been developed to ease the barrier of entry to 3D analysis, but to date most methods are generalized machine-learning applications that cannot separate background context from objects of interest, or custom-trained algorithms that require very large training sets (Lagree et al., 2021), (Stringer et al., 2021). Such enormous training data sets are not commonly available in either public or lab-curated micro-CT databases of specific millimeter to centimeter scale samples such as engineered materials, clinical biopsies, or human disease model organisms.

Apart from being a commonly studied model for multiple genetically and environmentally induced diseases, zebrafish (*Danio rerio)* models are used for micro-CT-based cellular and organ level analyses due to their size. Developing fish require millimeter-to-centimeter-scale fields of view to make it possible to image an entire animal in a single imaging session (Seo et al., 2015), (Babaei et al., 2016), (Hur et al., 2018), (Ding et al., 2019). Zebrafish have been used to generate genetic and environmentally induced models for clinically significant human blood diseases (Jin and Zong, 2011) which, alongside normal development, have been investigated through methods describing genetic expression, qualitative cellular morphology, and *in vivo* blood distribution (Brownlie et al., 1998; Liao et al., 2000; Paw et al., 2003; Shafizadeh et al., 2002), (Kemmler et al., 2023), (Varela et al., 2014)]. These advances, alongside future efforts to study zebrafish blood on a cellular level, would benefit from automated, quantitative characterization of a large proportion of the blood cells in a developing fish. Such an approach would add potentially meaningful detail to studies of normal and abnormal hematopoiesis, particularly in human blood disease models.

As a step towards computational phenotyping, we present an automated workflow, that involves (1) the imaging of whole zebrafish using histotomography, (2) detection and boundary definition of blood cells (defined as erythrocytes, lymphocytes and thrombocytes) in fish, (3) computation of volumetric and geometric features of segmented cells, and (4) derivation of statistically relevant changes caused by experimental variables (Fig. 1). Our approach provides a framework that processes and automatically describes the cellular morphology for individual blood cells of whole organism zebrafish histotomograms, the spatial distribution of blood cells across the entire organism, and a system for quantitative comparison between samples. The most biologically compelling results are a comparison between *Huli hutu* (*hht*) mutant zebrafish (mutant in *pola2*, DNA polymerase alpha B subunit) (Lin et al., 2021) and wild-type zebrafish, showing progressive degeneration of blood from 4-days-post-fertilization (dpf) to 5-dpf. The resulting contribution is a fully automated cell modeling workflow from 3D micron-scale micro-CT scans of whole organisms. We demonstrate the paradigm using modeling of blood cells in whole zebrafish, and the analysis pipeline can be potentially adapted to modeling other cell types in zebrafish and to modeling cells in other whole organisms.

**Figure 1:**
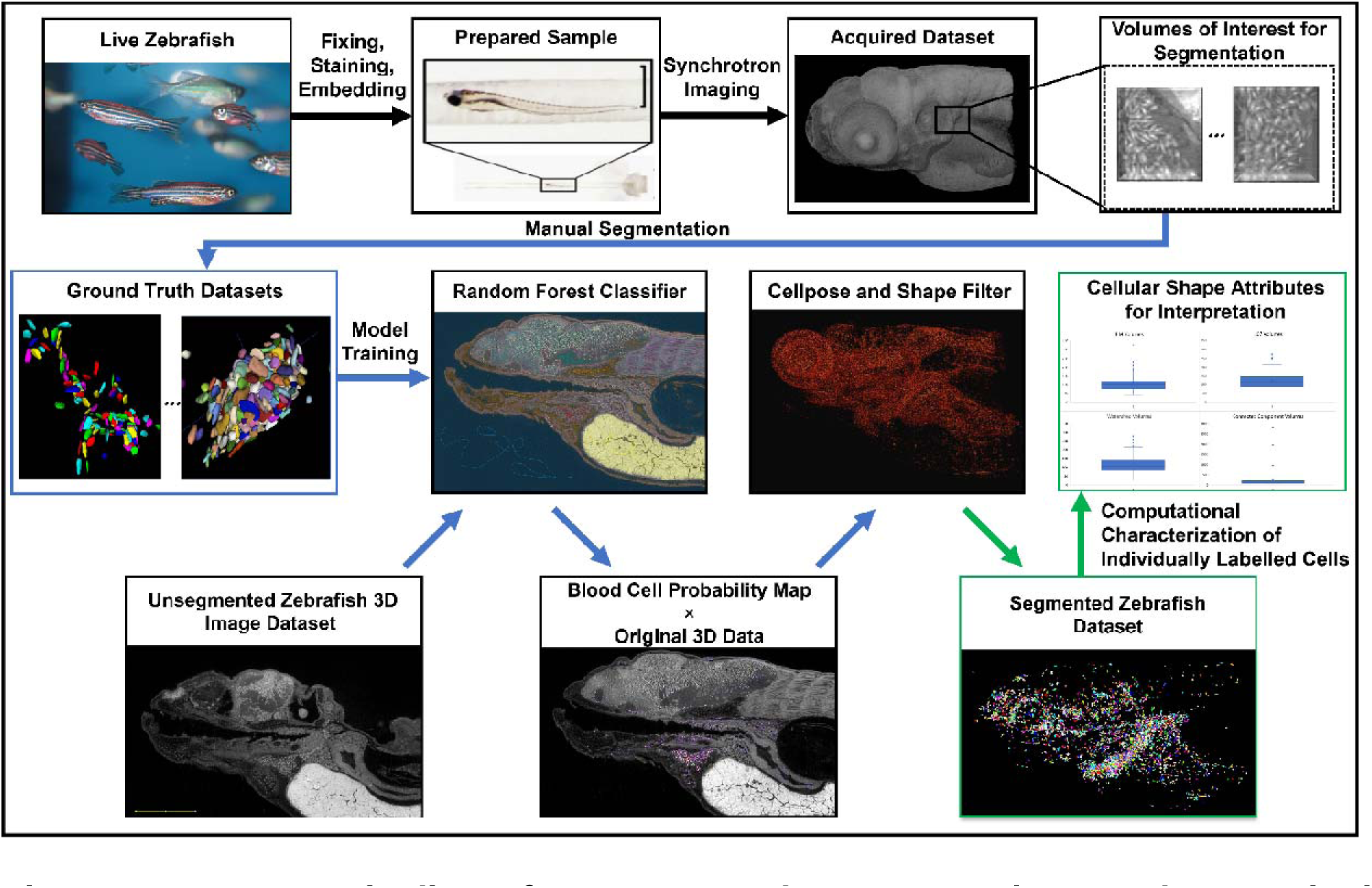
Data pipeline for automated segmentation and quantitative characterization. Samples are fixed, stained with PTA, and embedded in resin prior to scanning at synchrotron micro-CT facilities. Representative areas are selected for manual segmentation, enabling machine learning algorithm training and the establishment of ground truth validation sets. Whole unsegmented samples are processed through the automated segmentation pipeline and compared to the manually segmented volumes: cellular-level statistical shape and image intensity attributes are extracted from both sets and confirmed in the same regions. Processing multiple mutant and wild-type samples at different developmental stages allows for accurate blood cell comparisons between experimental and control groups.

## 2. Results

### 2.1. Identification of Blood Cells in Developing Fish

Blood cells in different stages of zebrafish development can be identified across multiple imaging modalities. To ensure that any quantitative characterization derived from micro-CT datasets reflects biological shape details identified from higher-resolution information, we compared zebrafish micro-CT data to corresponding histology slices and ssEM scans (Fig. 2, Fig. 3). 5-micron thick hematoxylin and eosin-stained histological sections are commonly used to study and identify zebrafish tissues and cells, including blood cells. We approximated these conditions by creating a “virtual section”: a maximum-intensity projection of the micro-CT data, taken through 5 microns of tissue, at a similar angle to an existing histology slice (Fig. 3 A, B). Blood cells look similar in both modalities, including shape, size, contrast, and the clear visible presence of both a cytoplasm and a nucleus (Fig. 3 C, D).

**Figure 2:**
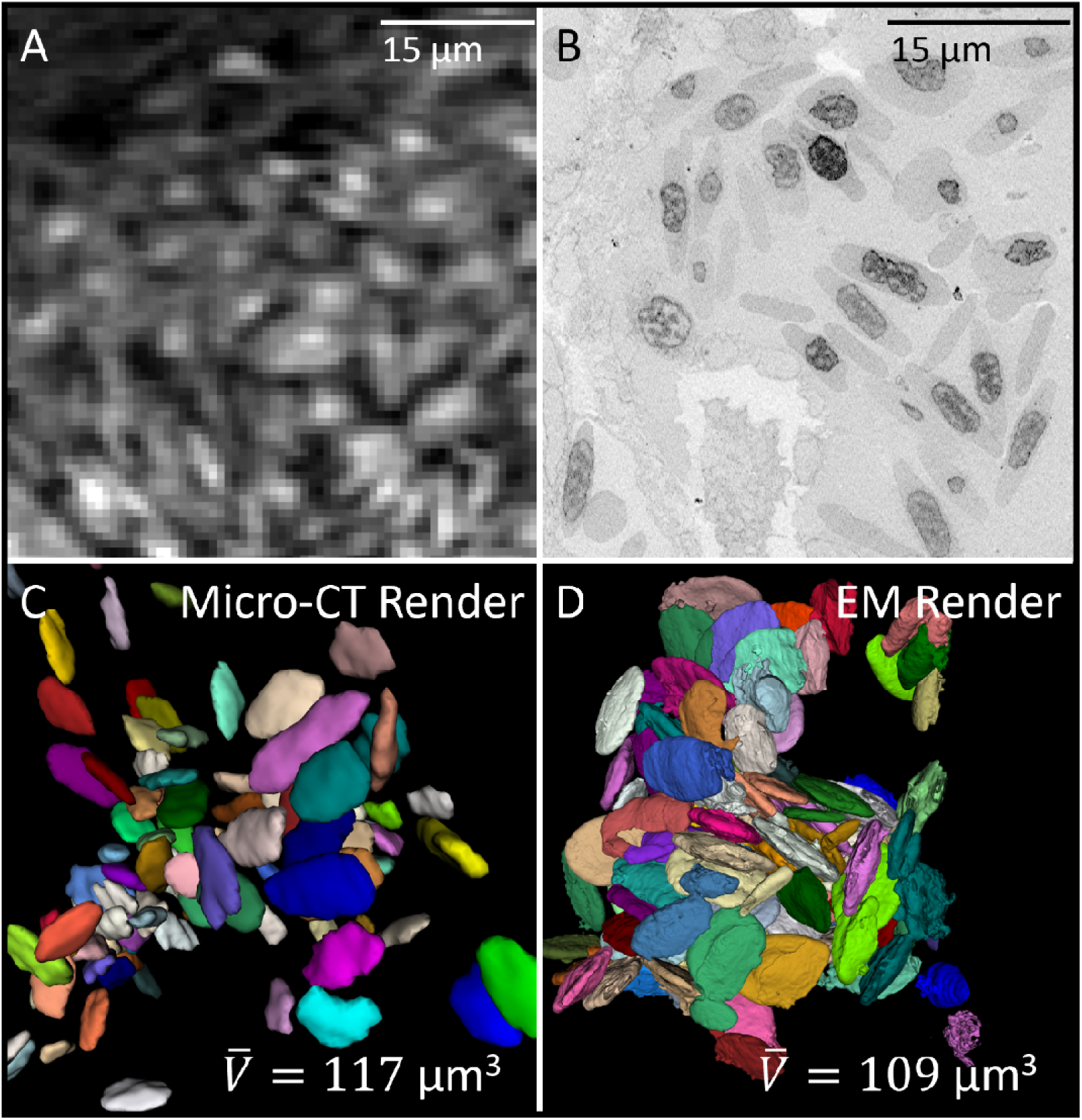
Comparison of slice images and 3D renders of blood cells acquired with micro-CT and serial-section electron microscopy (ssEM). (A) Blood cells from our 5-dpf dataset were compared to (B) a published SEM dataset of another 5-dpf fish. (C, D) Both datasets were automatically segmented using Ilastik and rendered in 3D. Average volumes of individual blood cells were compared between 3 manually segmented EM-scanned cells and the manually segmented micro-CT validation set from the heart, with the values differing by only 8 “µm3” (acquired using ITK-SNAP).

**Figure 3:**
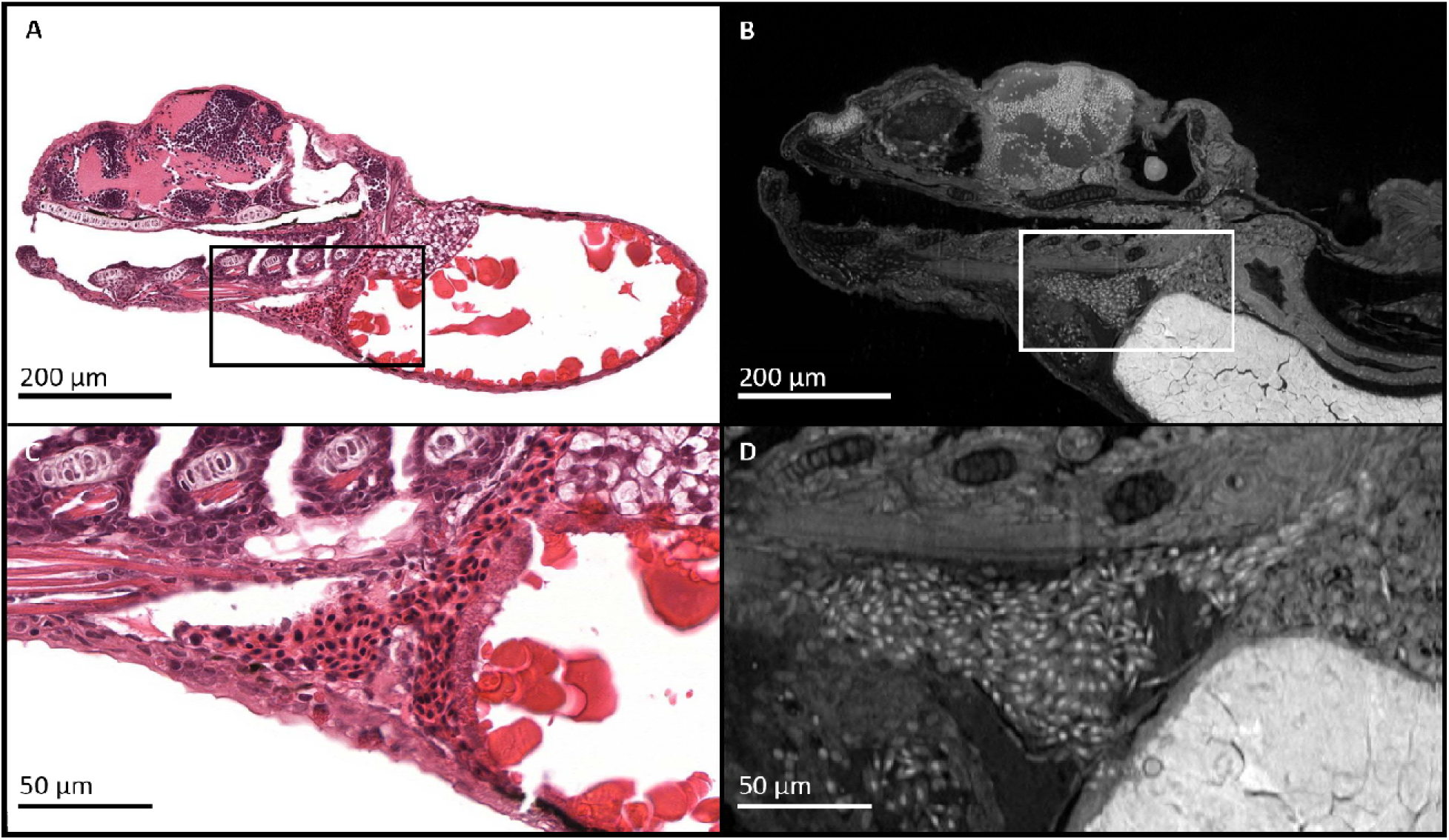
Blood cells were identified in micro-CT datasets based on their known staining patterns in histology. (A) Hematoxylin and eosin stained 5-micron thick slices of a normally developing 5-dpf zebrafish are compared to (B) a 5-micron thick maximum intensity projection taken at a similar angle from a micro-CT scan of a similar, normal 5 dpf fish. Staining patterns between histology and PTA-stained micro-CT images show similar staining across anatomic regions, including the heart (C, D). Individual nucleated blood cells can be distinguished across both imaging modalities for segmentation. Physical cutting artifact in the histology section push certain features out of the section (e.g. yolk in the bottom right of the zoom-in of the histology is partially missing due to loss during floating of the tissue section, and is fully present not present in the virtual section of the micro-CT dataset.)

### 2.2. Automated Segmentation Pipeline

Once blood cells were identified in micro-CT data, preliminary segmentation was done manually to characterize the shape statistics and distributions of normal cells. These validation sets were also used to both qualitatively and quantitatively test the degree of success of our automated segmentation pipeline. Automated segmentation of blood cells was divided into several major components. A random forest classifier was trained with manual segmentation data (unrelated to the validation sets) to generate a probability map of the blood cells within every sample. This result was then multiplied by the original data to decrease the signal of background tissue such as muscle, brain, yolk, and everything else that is not a part of a blood cell. A pre-trained neural network (Cellpose) that was adapted to this type of data was then fed the processed image to segment out areas of high probability. All potential blood cell segmentations were filtered by requirements of shape statistics for normal cells derived from the manually segmented validation sets.

Ilastik, an open-source random forest classifier was used to train a model for blood cell detection in developing zebrafish of various ages (Fig. 4). Datasets were divided into 9 classes (Fig. 4A) that encompassed a variety of tissue and cellular textures, improving the classification of blood cell volumes over multiple training steps that each added more data for the model to learn from (Fig. 4A’, A’’, A’’’). This model outputs a probability map of values of 0 through 1 for each voxel of the dataset, for each trained class. The probability map for blood cells has concentrated higher values in the heart, as well as areas containing other blood cells or tissue resembling blood cells (Fig. 4 B, B’). This probability map of the blood cells is then multiplied by the original data, reducing the signal in background tissue while retaining the original intensity values of the volumes of interest (Fig. 4. C, C’).

**Figure 4:**
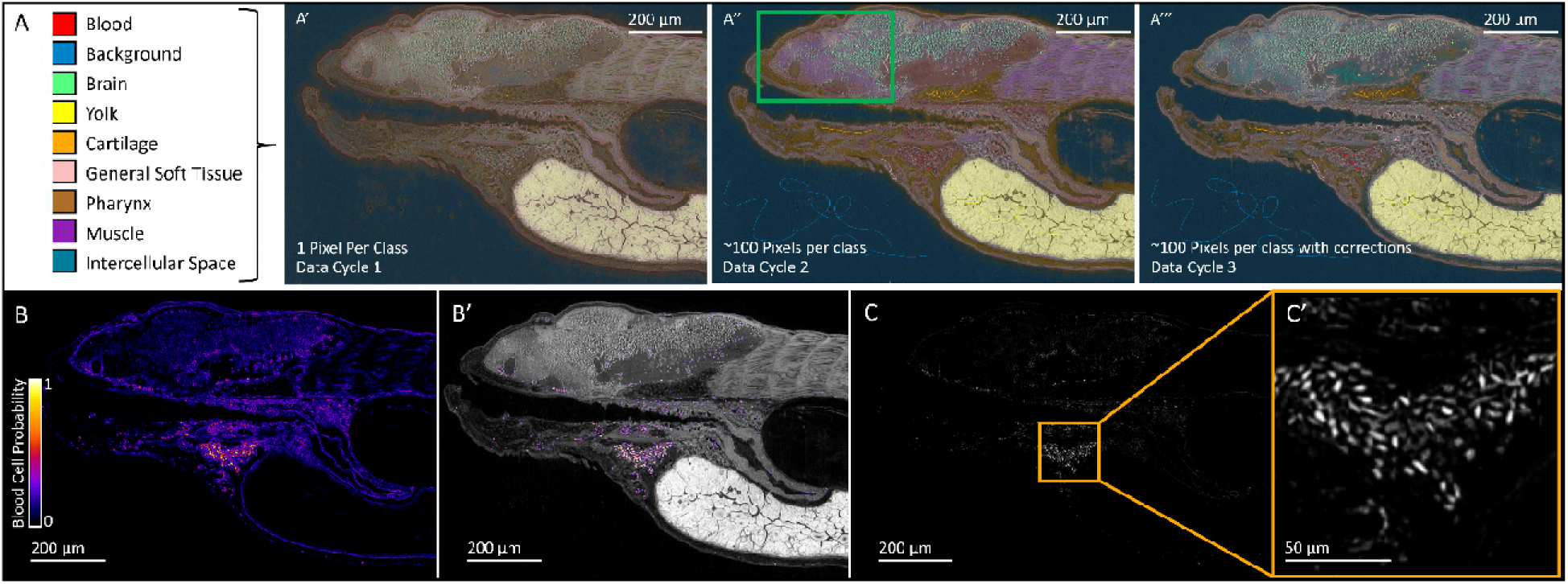
An iterative, machine learning-based training approach was used to isolate non-specific blood cell voxels for automated segmentation. (A) A 2D example of random forest classifier-based iterative model development is presented on a single slice of a wt 5 dpf fish dataset, where the pixel prediction of each tissue class improves with additional data. (A’) A single annotated pixel per class can separate yolk and empty background space but does not separate the tissues into distinct zones of confidence and is highly error prone. A round of basic annotation (A’’) significantly improves the result, but still shows classification issues in complex areas (e.g., brain regions interpreted as muscle in the green box). Additional annotation specific to those areas (A’’’) further refines the accuracy of the pixel classification. (B) Output from a trained Ilastik model is in the form of a probability map for each class. The probability map for blood cells in the same slice is shown as a heat map, and (B’) is overlaid back onto the original data. (C) Input to Cellpose for automated segmentation of individual cells was generated by multiplying the probability map by the original image, generating a dataset that retains any cellular properties in the volumes of interest ((C’) in this case intensity profiles of blood cells) while suppressing all other anatomic background.

The processed image is then fed through a cell segmentation neural net (Cellpose) to identify the boundaries between individual cells for preliminary segmentation (Fig. 5, 6). The preprocessing step (Fig. 5A, B) allowed us to utilize a highly accurate but generalized segmentation model for a very specific application on a single cell type, in what would otherwise be a very high-background dataset with various other tissues and cell types dominating the analysis. The preliminary segmentation results (Fig. 5C, D) include volumes of high probability, which are not always blood cells but are on occasion spaces composed of cells and tissues that resemble blood cell morphology (Fig. 5B, C, D). These 3D objects are filtered using shape analysis: computational measures such as volume, elongation, flatness, equivalent spherical radius, and principal axis lengths derived from the manually curated manually segmented validation sets were used to remove instances of false positives (Fig. 5E). The removal of false positives is most apparent in a 3D rendering of the results before and after filtering (Fig. 6). Post-filtering, blood cells are primarily retained in the expected regions, heart and dorsal aorta (Fig. 6A, B, Fig. 9).

**Figure 5:**
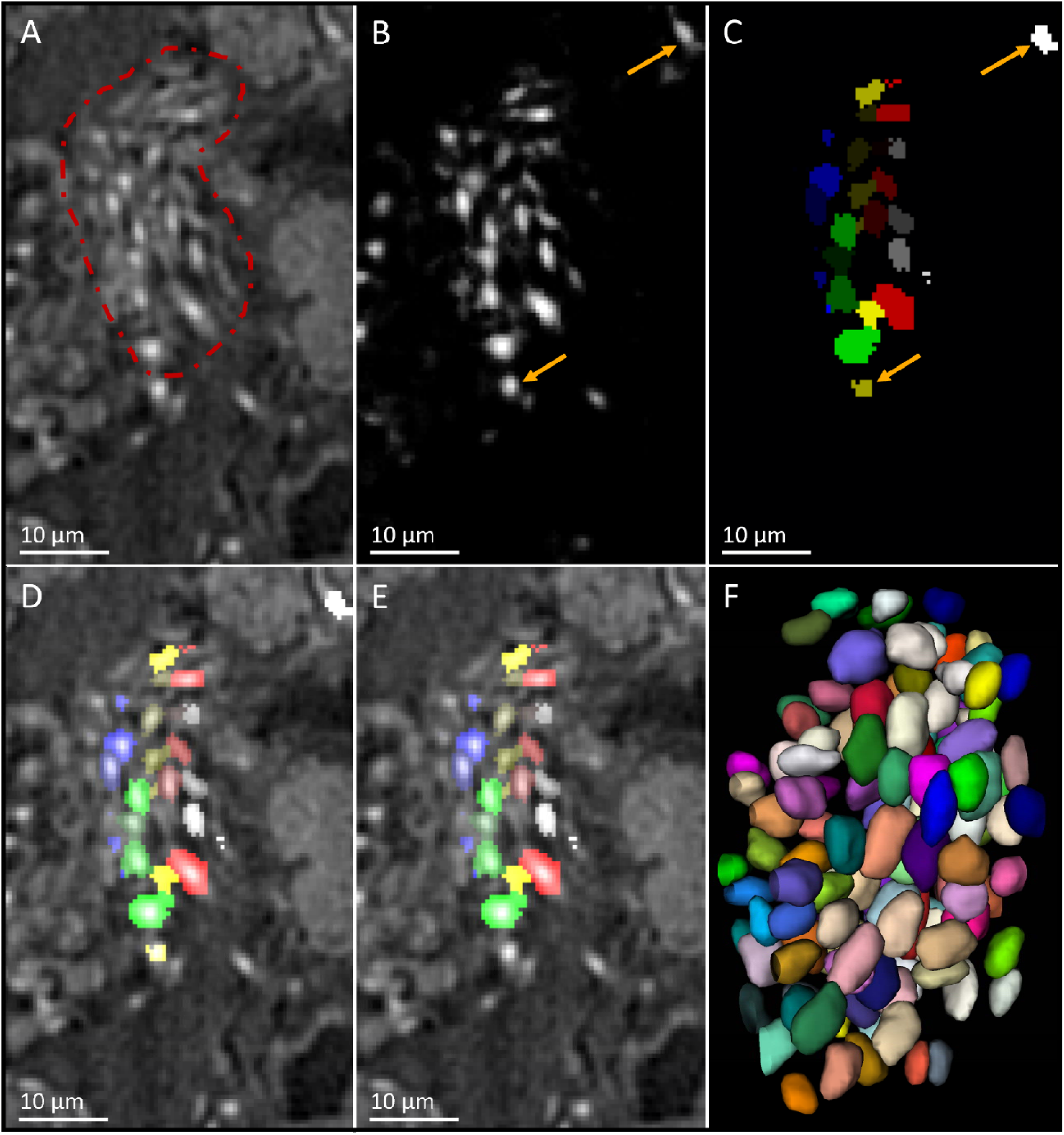
Automated segmentation workflow visually confirms boundary detection of cells and is further improved by shape statistic filtering. (A) A 2D slice of micro-CT data from a 5 dpf fish is shown with the heart highlighted in red. (B) The same region is presented as a result of the image multiplied by the corresponding probability map. Arrows point to regions of high probability that resemble blood cells but do not share the manually-derived blood cell shape statistics. (C) Initial segmentation results identify these areas, despite (D) visually confirmed accurate boundary detection. (E) Automated removal of any detected objects that do not fall within the known shape parameters of zebrafish blood cells removes these objects. The 3D render of segmented blood cells in entire region is shown in (F).

**Figure 6:**
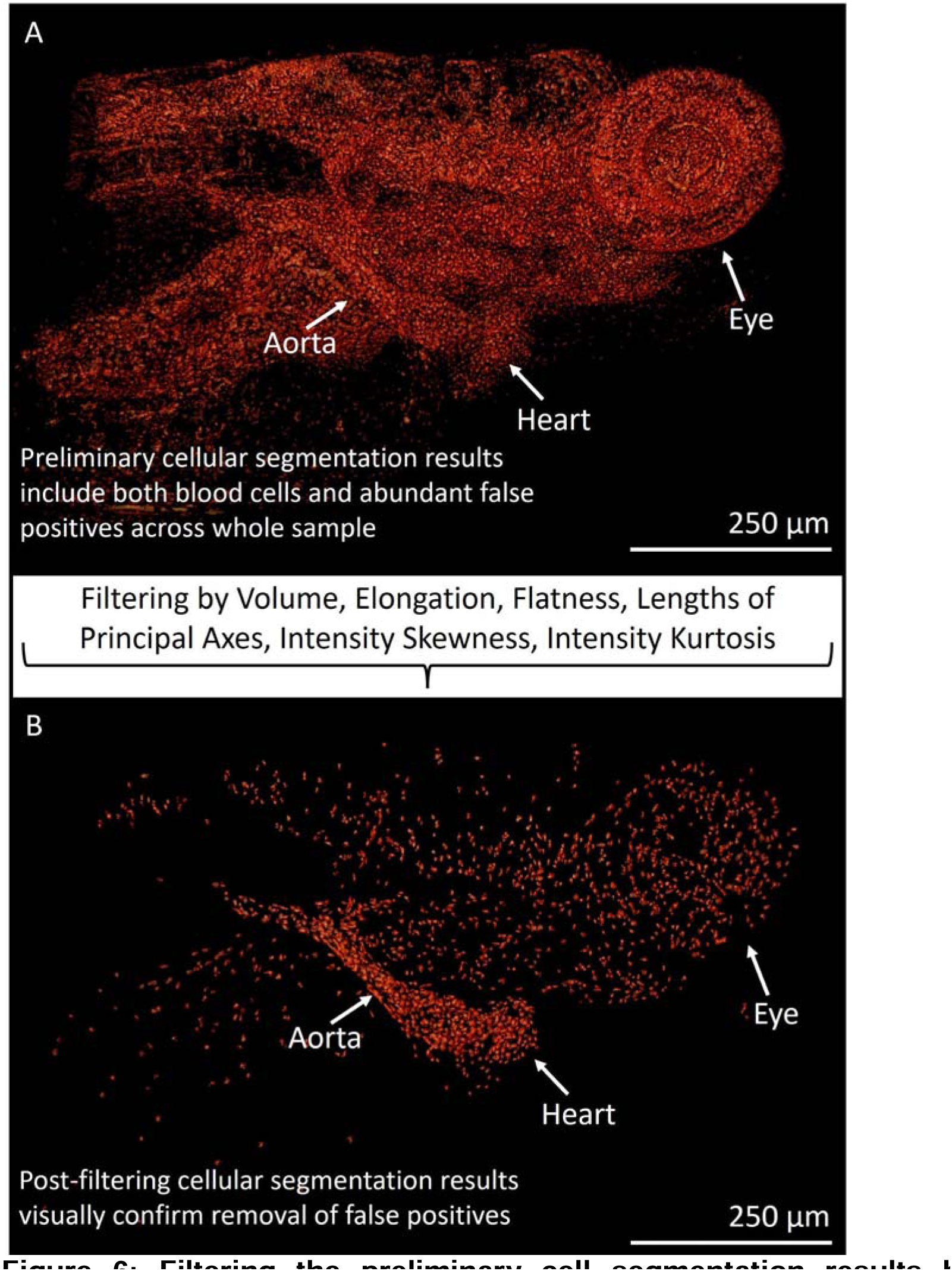
Filtering the preliminary cell segmentation results by biologically informed shape and intensity statistics removes false positives from the dataset while retaining segmented blood cells in expected regions. (A) Cellpose output of the processed Ilastik data filters cells from most areas of the body but leaves false positives in tissues that resemble blood cells in texture, such as neural nuclei in the brain and eye. (B) Filtering these preliminary results by known biological characteristics of blood cells, obtained from automated measurements of manually segmented validation sets, removes these false positives while retaining true positives within expected regions such as the heart and major vessels.

### 2.3. Segmentation Optimization and Validation

The accuracy and precision of the automated segmentation results vary depending on parameters set during the formation of the pipeline and during data acquisition, and thus an F1 score-based method of validating various test results was used. This allowed us to compare different automated segmentation outputs with manually segmented validation sets to maximize the quality of segmentation for batch processing (Fig. 7).

**Figure 7:**
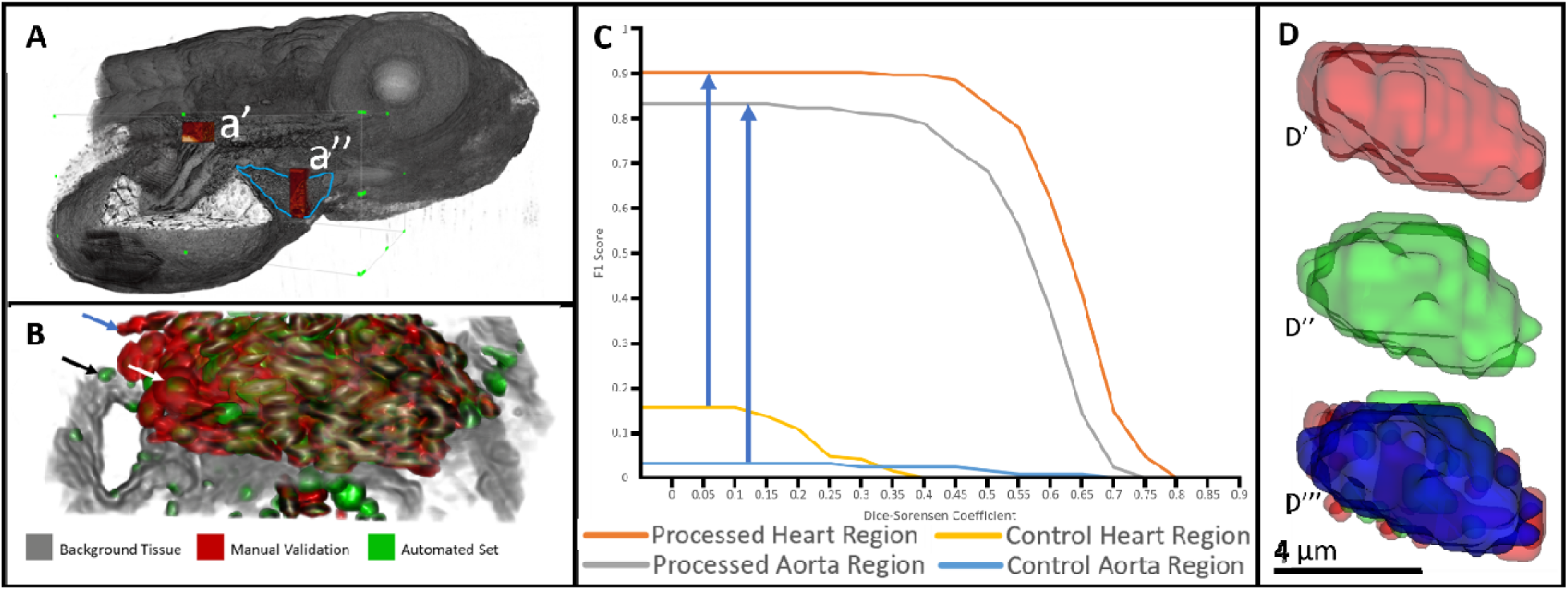
Degree of physical overlap, measured as an F1 score, between automatically segmented blood cells and manual sets validates the accuracy of our automated workflow. (A) Validation regions in 5dpf fish were taken from a section of the dorsal aorta with few blood cells and abundant tissue background (a’), and the heart, which is dense in blood cells and low in other biological background (a’’). This allowed us to test automated segmentation accuracy in widely varied regions. (B) A region used for validation is shown with various degree of overlap between manually and automatically segmented cells. Arrows point to examples of clear false positives (black arrow), clear false negatives (blue arrow) and potential true positives (white arrow). (C) Optimizing similarity of automatically segmented cells to manual validation sets was done quantitatively. F1 scores were generated using increasing Dice-Sorensen coefficients as requirements for what degree of overlap between a manually segmented and automatically segmented cell qualifies as a true positive. Pre-processing the raw micro-CT data using the probability map and post-processing the data by filtering cells using the known biological shape statistics of blood cells increased the performance of the automated segmentation pipeline, both in validation regions of the heart and aorta (blue arrows). (D) The overlap (D’’’) of a single cell, segmented manually (D’) and automatically (D’’), is shown as a 3D render. Qualitative observation of an optimized output shows a large degree of overlap (blue) between the segmented data.

Volumes for validation sets contained several hundred cells and were taken from the heart and dorsal aorta (Fig. 7A) to include signal heavy and background heavy data, respectively. Because our criteria for considering true positives involves not only detection of any individual cell in the same area as in the validation set, but also shape and volume characteristics, calculated using the Dice-Sorensen coefficient for each corresponding cell in the automatically- and manually-segmented sets. Overlap between individual cells in the automated and manual sets vary (Fig. 7B), and criteria for what is considered a true positive can change the F1 score significantly. Using a discrete cutoff value for what is considered a true positive and segregating all other instances into false positives and false negatives, an F1 score was generated for each integer cutoff ranging from 0 to 1 (Fig. 7C).

Optimizing the pipeline (Fig. 8) and performing all the preprocessing steps described ensured a ∼0.83-0.92 F1 score in both validation regions of the 5-dpf wt fish with little drop-off until a Dice-Sorensen cutoff of 0.5 (Fig. 7C). Optimization was performed for each variable in the pipeline, covering a range of values that increased, maximized, and then decreased F1 score (Fig. 8A, B). Variables optimized included filtered volume, elongation, flatness, principal axes lengths, intensity skewedness, and intensity kurtosis. After optimization of the pipeline, F1 score plots were compared to those generated by applying the Cellpose model to the raw, unprocessed micro-CT data in the same regions. Neither of these control groups generate an F1 score over 0.17 for either validation set comparison (Fig. 7C). A 3D render of the overlap between a single manually and automatically segmented, post-optimization cell is shown (Fig. 7D) in comparison to under and over segmented results. The renders visually confirm that the F1 score optimization is performing as expected on a cellular level; this step makes the automatically segmented cells better reflect the boundaries and shapes of the manually segmented cells (Fig. 8B).

**Figure 8:**
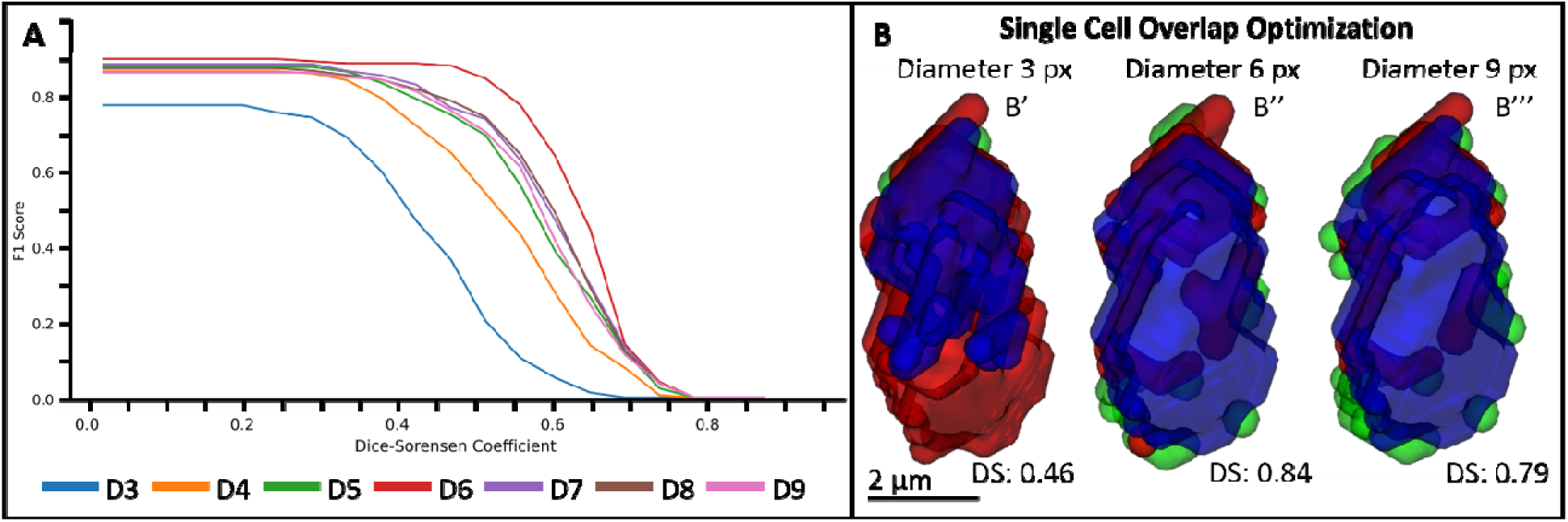
Automated segmentation pipeline was optimized through maximizing Dice-Sorensen coefficients between automated results and manually segmented validation sets. (A) Generated F1 scores for various Dice-Sorensen cutoffs are shown for cells automatically segmented in the heart (a” in Figure 7). The pipeline was adjusted to segment cells with a series of average diameters of 3 pixels (D3) through 9 pixels (D9). The closest computational similarity between automatically and manually segmented cells was obtained for an average diameter of 6 pixels, which corresponds to the average spherical radius of manually-segmented zebrafish blood cells. Visual assessment for an individual cell in the center of the distribution illustrates this result (B). The same cell is rendered as the manual segmentation from the validation set (red), automated results (green), and volumes of overlap (blue) for diameters of 3 pixels (B’), 6 pixels (B’’), and 9 pixels (B’’’). Visual results confirm the highest Dice-Sorensen score when using an expected diameter of 6 pixels.

Our computational validation was confirmed by qualitative assessment of blood cell distribution within the sample (Fig 9). Blood cells, while distributed throughout the body, generally do not appear in predictable tissues lacking direct access by blood vessels at 5-dpf in development such as the swim bladder muscle walls, brain regions occupied by neural nuclei, and the yolk sac (Fig. 9B-E). The largest concentration of blood cells is located within the heart and dorsal aorta, as well as various extending vessels appearing as lines of cells (Fig. 9A).

**Figure 9:**
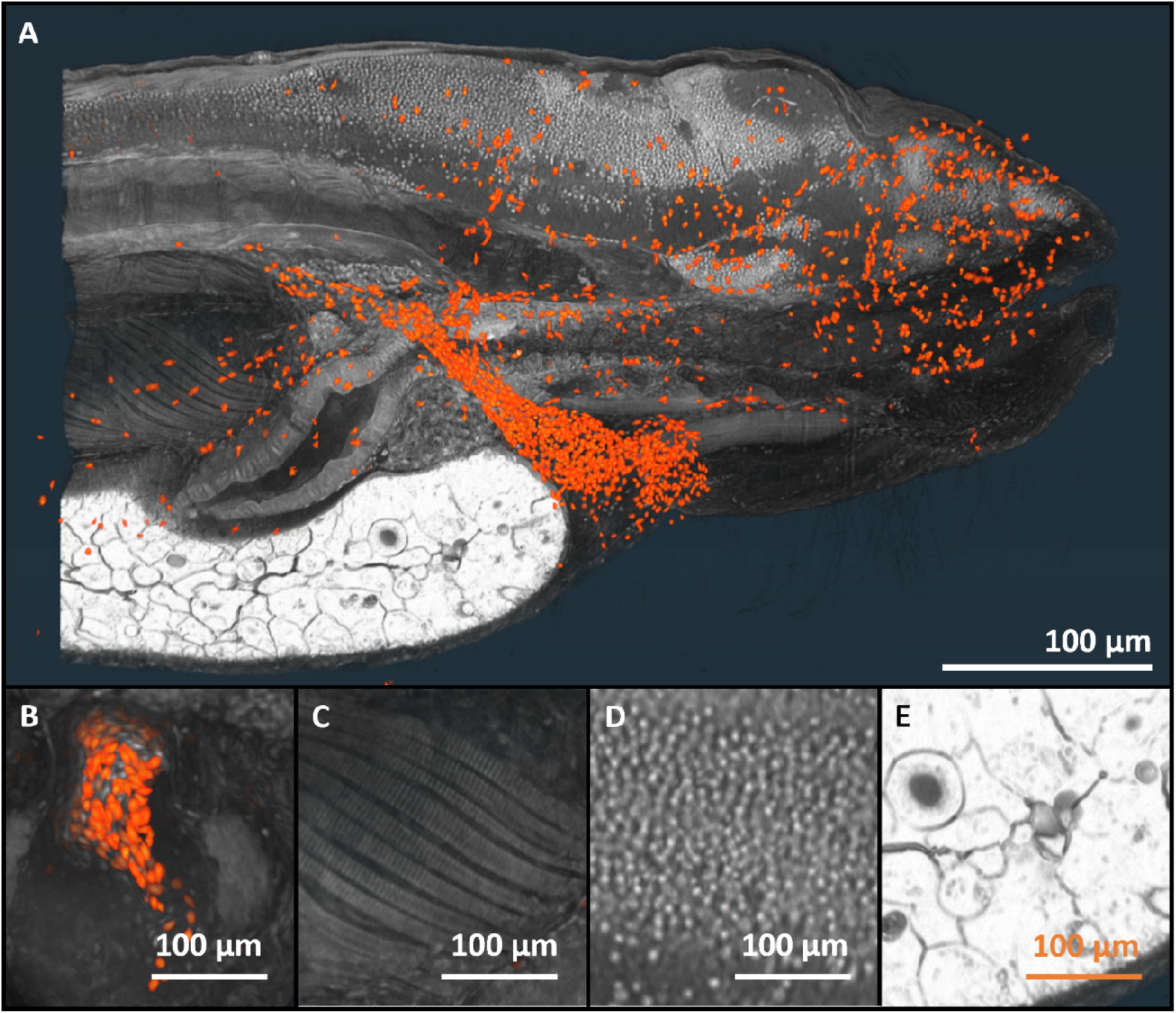
Rendering of detected blood cells in context of original data. (A) All detected blood cells in a 5-dpf zebrafish are shown overlaid onto a rendering of half of the background anatomical data, cut sagittally. (B) A transversely cut, zoomed-in render of the detected cells in the heart region of the 5-dpf fish is shown, illustrating a high concentration of blood cells. A zoomed-in render of the raw zebrafish data with all detected cells is provided across large areas of the (C) air bladder muscles, (D) brain neural nuclei, and (E) yolk sack to emphasize the absence of detected blood cells in regions expected to lack them.

These vessels were separately investigated for cellular detection which was confirmed in vessels running from the dorsal aorta into the eyes, brain, and adjacent to the yolk (Fig. 10). An advantage of isotropic 3D data is the ability to visualize particularly tortuous vessels (Fig. 10B) as straight tubes taken across one plane for quick visual confirmation of blood cell detection (Fig. 10C). Taken together, the detection and distribution of blood in expected structures, alongside the quantitative assessment of our automated segmentation method, illustrates the accuracy with which the automated algorithm can extract cells for shape characterization and further study via comparison to other similarly processed samples.

**Figure 10:**
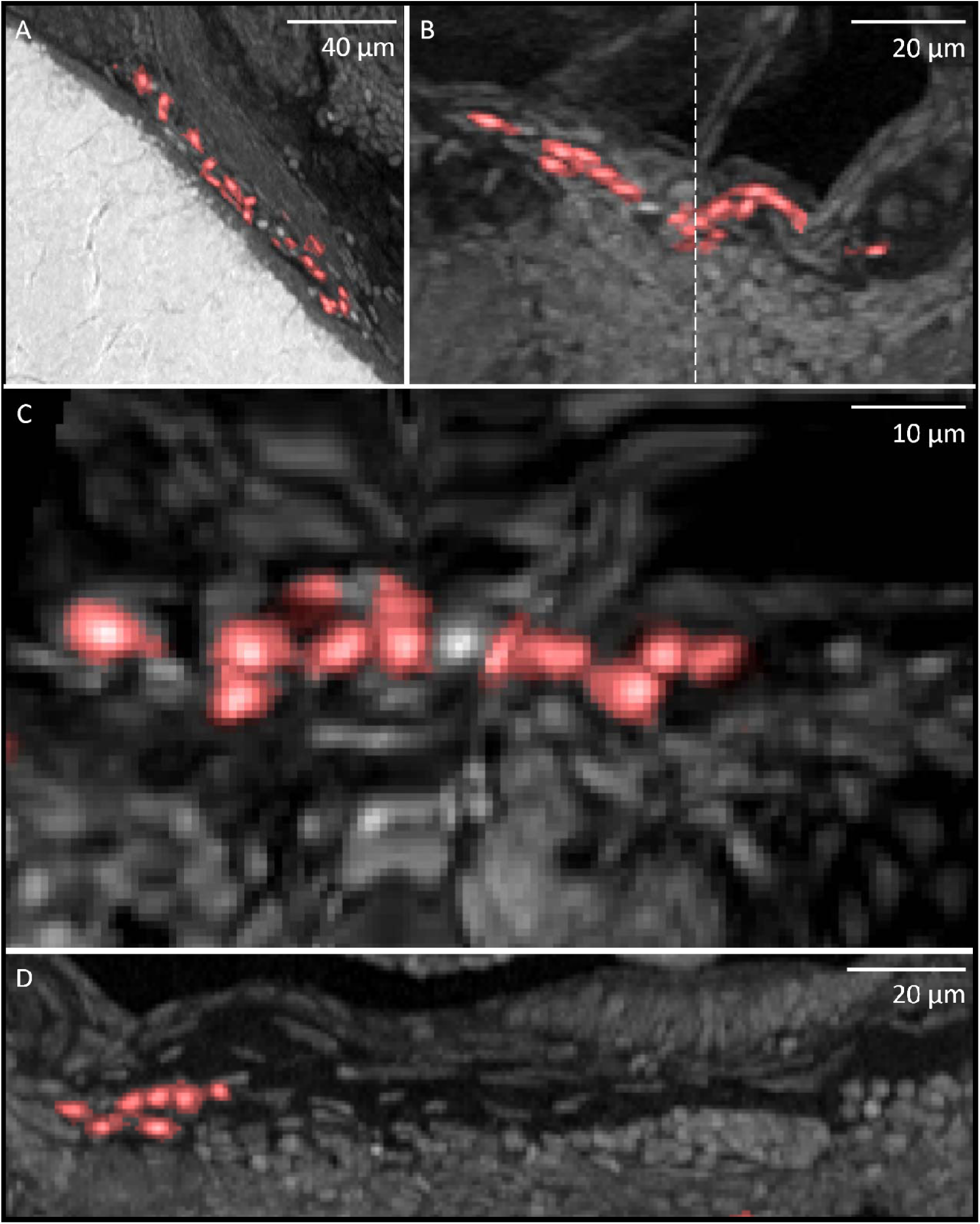
The automated segmentation pipeline detects blood cells in thin blood vessels. (A) A 5 µm thick MIP of a vessel approximately 10 µm in diameter running adjacent to the yolk shows blood cells in various orientations. (B) A MIP of a thinner, approximately 5 µm thick blood vessel running to the brain is presented with overlaid detected cells oriented in the same direction. A dotted line is shown between its two halves where the tortuous nature of the vessel travels out of slice. (C) The nature of 3D data allows the same vessel in (B) to be straightened out digitally for interrogation as a single line. (D) The walls of thin vessels with few blood cells are easier to visualize due to their high amount of empty space.

### 2.4. Characterization of Wild-Type Development and Comparison to *hht*

Our analysis focuses on the commonly studied stages of development of 4-dpf and 5-dpf, between normal wild-type growth and the degradation of the *hht* mutant. From our knowledge of striking differences in histological appearance between wild-type and mutant fish, we expect a large statistical difference between these two genetic backgrounds. Strong differences need to be seen in quantitative shape statistics in this setting if we are to be convinced that our approach can be useful for studying disease models. Because the *hht* mutant lacks a functional DNA polymerase *pola2*, it relies on its initial amount of maternally provided polymerase Lin et al. . This supply rapidly degrades, resulting in the fish starting development normally but quickly deteriorating on a cellular and organ level due to its inability to perform normal cell division during a time of growth. This is reflected in our findings, as the number of blood cells between wild-type 4- and 5-dpf fish quadruple in number (from 1057 to 4491), while the 5-dpf *hht* sample contains approximately the same number of cells as its 4-dpf *hht* counterpart (from 714 to 882) [Tables 1-4].

The difference in development between the genetic backgrounds is further observed in the corresponding blood cell shape statistics between the *hht* and wild-type datasets (Fig. 11). Visualization of the shape attributes reveals significant differences across sample groups, as genotypes are readily discernible by plotting cell volumes in a histogram (Fig. 11A). Wild-type cells have a significantly larger mean volume than *hht* mutant cells, even when controlling for age. Within genotype, wild-type cells increase in volume and quantity from 4- to 5-dpf age, supporting prior biological knowledge of zebrafish development (Ransom et al., 1996) (Fig. 11C). *hht* cells yield a different distribution of volume from 4- to 5-dpf, exhibiting a notably smaller mean volume and no clear increase in quantity. Further analysis of all shape statistics extracted through the pipeline provides a framework for further machine learning and statistical inference. Dimensionality reduction performed using linear discriminant analysis demonstrates the ability of the pipeline to segregate different cell phenotypes based on age and genetic background (Fig. 11B). Cells from each of the 4 groups analyzed in this pipeline are readily discernible when the data are plotted across the first 3 linear discriminants. Application of semi supervised learning approaches to the data yielded by this pipeline provide proof of concept in advancement towards computational phenomics, demonstrating the ability to separate segmentation results by age and genotype.

**Figure 11:**
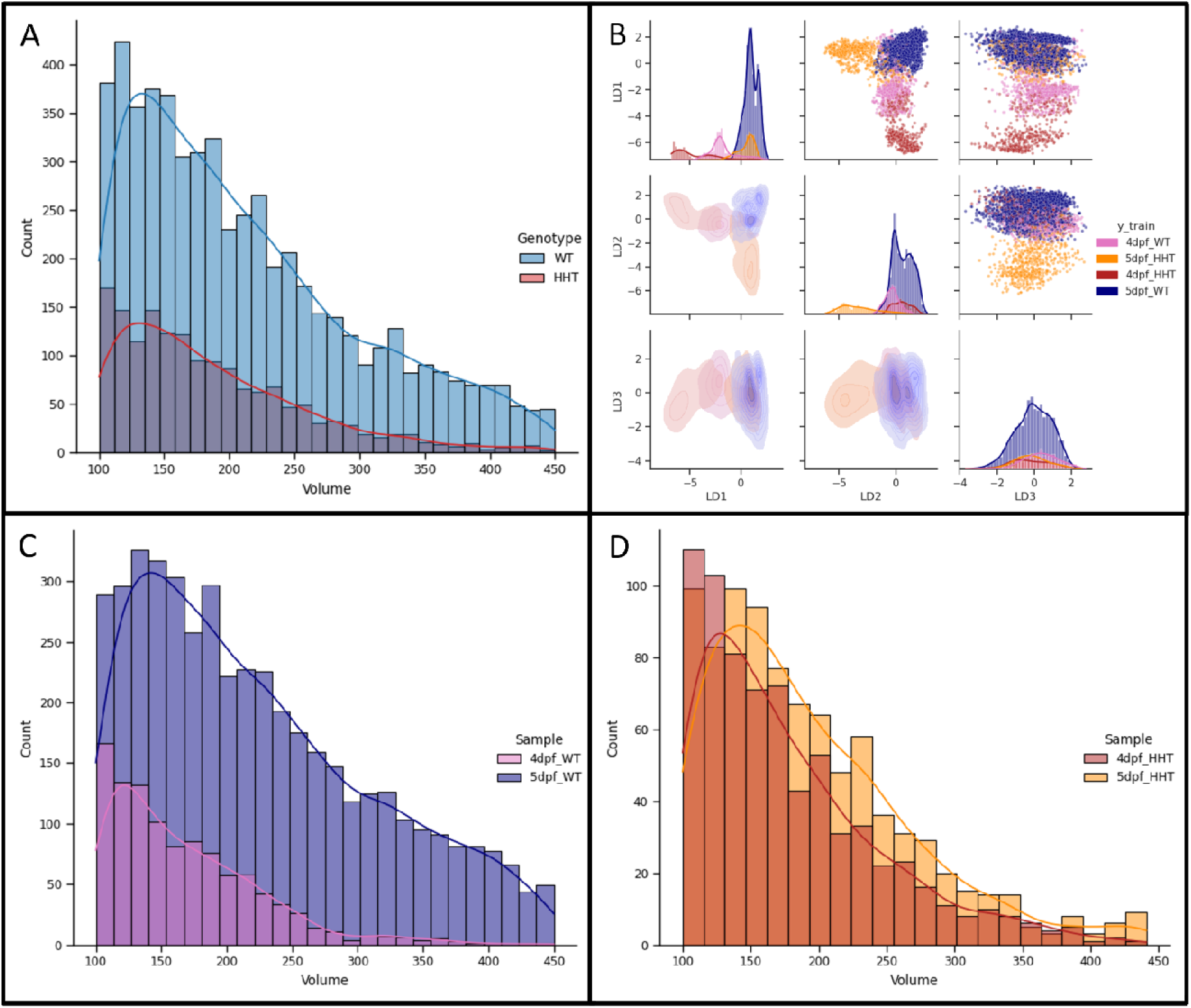
Comparison of blood cell shape statistics reveals distribution differences between samples. (A) Distributions of wild-type (WT) and *hht* blood cell volume measurements extracted from the automated segmentation pipeline are shown plotted according to zebrafish sample. (B) Linear discriminant (LD) analysis applied to shape attributes from each sample reveals separation of cells by genetic and age phenotypes. (C) Distribution of cell volume within the wild-type genotype is shown stratified by age. (D) Distribution of cell volume within the *hht* genotype is shown stratified by age.

These differences between the four genetic and development conditions can also be easily observed qualitatively (Fig. 12) in terms of blood cell distributions and gross anatomy. The wild-type and *hht* samples take on distinct body shapes, which diverge from the wild-type not only in cellular features but in organ size, shape, and distribution. The heart is pronounced and takes up much more space in the wild-type samples, compared to the underdeveloped and rapidly dying organ present in the *hht* fish that holds the highest concentration of blood cells (Fig. 12A-D). The shape and size differences are also apparent when the average principal axes lengths are used to approximate the average blood cell from each genetic background as an ellipsoid.

**Figure 12:**
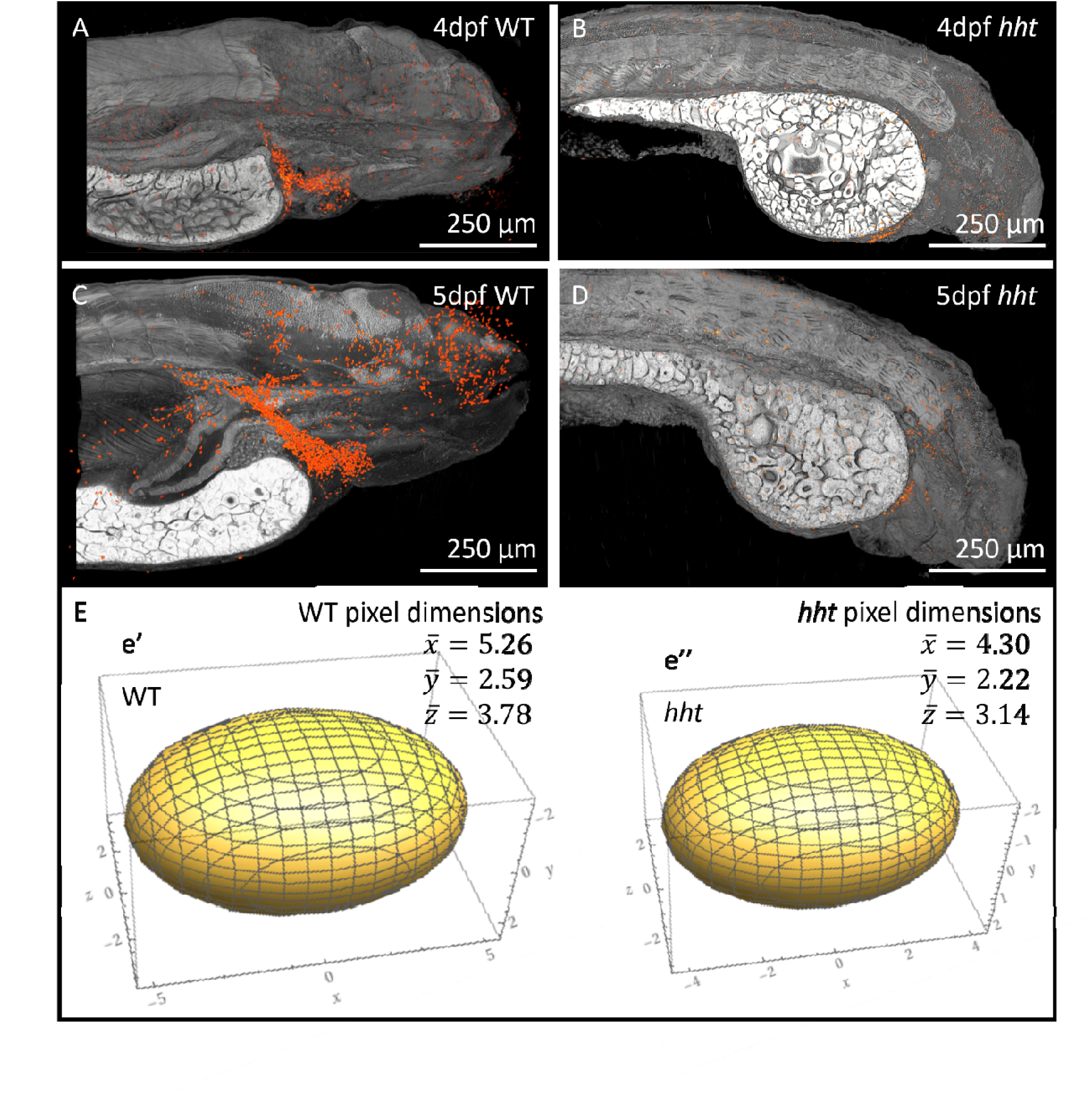
3D renders of blood cell distributions in 4- and 5-dpf wt and *hht* mutant fish. Blood cells (rendered in red) are shown overlaid onto a cut-in render of the background datasets. Cellular distribution is visibly different across the widely varied shapes of the WT and *hht* samples. (A, C) The highest concentration of blood cells in the 4- and 5-dpf WT fish can be seen in the heart and dorsal aorta, with various blood cells distributed through vessels in the body. (B, D) This distribution is roughly conserved in the *hht* samples, though the progressive degradation of the cells and organs alters the heart shape, cell counts, and (E) cell shape between the 5-dpf samples.

## 3. Discussion

### 3.1. Model Confirmation and Blood Cell Development in Zebrafish

As the interrogative power of imaging techniques increases both data quality and quantity for disease modelling, developmental studies, and clinical diagnosis, the need for automated analytical methods capable of processing 3D image datasets also increases. Here, we show the development and application of a segmentation and analysis pipeline that accurately characterizes and compares blood cells in micro-CT scans of developing zebrafish through an informed understanding of biologically constrained shape attributes. Our method accurately segmented blood cells from two distinct developmental stages and genetic backgrounds, extracted shape-defining statistics from all samples, and then was able to show differences between the datasets sufficient to classify new, unknown blood cells into these groups based on those same attributes.

The segmentation and analysis of blood cells distributed across the whole-organism offers a set of unique advantages by providing cell count, cell characterization, and anatomical background for each sample analyzed. The difference in the distributions of the cellular attributes between the genetic backgrounds is supported by our finding that total cell count was increased between 4- and 5-dpf wild-type fish but not between 4- and 5-dpf *hht* samples, corresponding with what is known regarding *hht* degeneration during development; as the amount of pola2 decreases in the mutant and cells lose the ability to replicate, the number of cells and size of the fish should stop increasing and eventually cause death around 14-dpf. The visualizations of the original scanned fish data overlaid with the segmented blood cell data confirms this difference in anatomical shape and organ distribution and allows researchers to explore the full depth of both the masked cells as well as any background tissue of interest. Cellular counts, statistical shape analyses, and distribution of cells across the whole organism provides both a quantitative and qualitative method to confirm the presence of cells of interest in expected tissues, offers a comparison to samples from different experimental groups, and allows for discovery of cell distributions in unexpected locations.

We anticipate that the most direct practical application for this approach is the confirmation of new and established zebrafish blood disease models. These include already generated mutations and environmentally induced phenotypes simulating conditions such as sickle cell anemia, hemolytic anemia, congenital dyserythropoietic anemia type II, hereditary spherocytosis, and congenital sideroblastic anemia (Shafizadeh et al., 2002), (Paw et al., 2003), (Liao et al., 2000), (Brownlie et al., 1998), (Williams-Pate & Gahr, 2019). Each of these models may benefit from showing quantitative cytological changes in red blood cells alongside anatomical background. This analysis, presented as is with no alteration, can be directly applied to such models to answer questions regarding the degree of change in blood cells compared to normal development while confirming that other cell types and organ systems remain unchanged or are altered as expected. This is accomplished by analyzing datasets with cell populations in the thousands, without resorting to extensive manual segmentation work. Additionally, studies of other fish models such as teleost, carp, and tilapia could benefit from this analysis pipeline as their blood cells are also nucleated and resemble the zebrafish in shape, and size (Megarani et al., 2020). Taken together, our analysis pipeline and results represent a step towards comprehensive 3D computational phenotyping in whole vertebrate systems on a cytological scale.

### 3.2. Generalizability of Segmentation Method for Sparse Datasets

While the specific application of our segmentation method was focused on characterizing and comparing the blood cell populations of developing zebrafish, this study in conjunction with previous work (Ding 2019) shows flexibility of the general approach to more than a single cell and/or tissue type. The use of generalized and random forest classifier-based steps in the segmentation method (Ilastik, Cellpose) enables the same type of approach on other micro-CT datasets seeking to segment cellular and organ structures but lacking large training datasets. Random forest classifier frameworks can provide accurate classification-based masking from sparse training data, a necessary condition for the processing of most micro-CT datasets specific to a single organism and a single cell type. Open-source applications such as Ilastik allow for the generation of probability maps of as many classes of cells and/or organs as necessary, allowing for the training of all segmentation targets of interest in parallel on the same dataset.

Masking out only the volumes containing the cells of interest via probability maps allows generalized approaches such as Cellpose to be used on each class, one at a time, focusing on the segmentation of cells within the pre-segmented data without contending with the biological background of the rest of the organism. Filtering preliminary results by shape statistics informed via established biological constraints limits the number of false positives within the detected objects. Finally, a comparison to manual segmentations established as ground truth within selected volumes of samples ensures accuracy of the approach and can inform the user of the requirement for a more robust training set. It is our hope that the generalizability of this approach will provide a pathway for extending it to other cell types within zebrafish and other organismal scans at different stages of development, and to clinical samples.

### 3.3. Future Work

The analysis framework applied to blood cells here can be extended to other biological imaging datasets and cell types. The detection and characterization of other cells, tissues, and model organisms will require customizations such as the selection of classes for segmentation and choice of shape statistics to measure and filter by. Additional procedures such as preliminary organ segmentation, using the same presented method on lower resolution images, can be implemented to restrict analysis to volumes of interest. Cell types with less defined borders than blood cells will pose a greater challenge to accurate segmentation. The ability to differentiate cell boundaries and cell types can be expected to be improved by increased resolution such as that which has recently become available using new micro-CT imaging systems (Yakovlev 2022). This work with blood cells in developing zebrafish provides a framework for commonly studied disease models that may benefit from automated segmentation and shape-based characterization of cells in their anatomically relevant context. Increasing accessibility to histotomographic imaging would allow computational phenomic approaches to evolve across model systems.

## 4. Materials and Methods

### 4.1. Key Resources Table

| Reagent type (species) or resource | Designation | Source or reference | Identifiers | Additional information |
| --- | --- | --- | --- | --- |
| Strain (*Danio rerio*) | Wild-type Ekkwill | ZFIN ID: ZDB-GENO-990520-2 |  |  |
| Genetic reagent (*Danio rerio*) | *huli hutu* | Mohideen et al., 2003 |  |  |
| Software, algorithm | Avizo | Thermo Fisher Scientific | SCR_014431 | version 2021 |
| Software, algorithm | Fiji/ImageJ2 | https://fiji.sc/ | SCR_002285 |  |
| Software, algorithm | Ilastik | Sommer et al., 2011 (http://ilastik.org/) | SCR_015246 | version 1.3.3 |
| Software, algorithm | ITK-SNAP | Yushkevich et al., 2016 (http://www.itksnap.org) | SCR_002010 | version 3.4 |
| Software, algorithm | OpenSeaDragon | https://openseadragon.github.io/ |  |  |
| Software, algorithm | TomoPy | Argonne National Labs (http://tomopy.readthedocs.io) | SCR_021359 |  |
| Software, algorithm | SimpleITK | Yaniv et al., 2018<br>Beare et al., 2018 (https://simpleitk.org/) | SCR_001149 |  |
| Software, algorithm | Cellpose | Stringer et al., 2021 (https://www.cellpose.org/) | SCR_021716 | version 1.0.2 |

### 4.2 Zebrafish Husbandry and Sample Preparation

Zebrafish were housed and raised in the Penn State Zebrafish Functional Genomics Core, which is equipped with 2 recirculating aquaria with 14:10 hour light:dark cycle and average temperature of 28°C. The wild-type zebrafish (Ekkwill strain) and *hht* mutant (Mohideen et al. 2003) were fed three times a day with diet consisting of brine shrimp and flake food. The *hht* mutants were maintained as heterozygous because the nature of the mutation is recessive larval-lethal. Zebrafish were set up for mating using Aquatic Habitat breeding tanks with dividers the afternoon prior to embryo collection. Once collected, the embryos were disinfected in 10% Ovadine (Syndel) for 2 min at room temperature followed by three washes with charcoal-filtered water. The embryos were incubated in 28°C incubator to maintain consistent development. Developmental staging series of fish were done based on Kimmel et al. (1995). Homozygous *hht* mutants were identified through gross phenotypes, including dorsally curved tail, small head and eyes, and enlarged yolk under stereomicroscopes at 2-dpf.

Larval (2-, 3-, 4- and 5-dpf) zebrafish were euthanized in ice-buffered MS-222 (400 mg/L) solution (Argent Chemical Laboratories, Redmond, WA). The fish were fixed in pre-chilled 10% neutral buffered formalin (NBF) (Fisher Scientific, Allentown, PA) overnight at room temperature (Lin et al. 2018). The fish were stained with 0.3% phosphotungstic acid (PTA) buffered in 100% ethanol for 24 hr at room temperature on a rotator and embedded as described in Lin et al. (2018). All procedures on live animals were approved by the Institutional Animal Care and Use Committee (IACUC) at the Pennsylvania State University, ID: PRAMS201445659, Groundwork for a Synchrotron MicroCT Imaging Resource for Biology (SMIRB).

### 4.3 Micro-CT Image Reconstruction and Visualization

All stained samples were imaged at the Argonne National Laboratory synchrotron facilities during two visits in 2011 and 2013 and reconstructed on site. Each individual scan was acquired as 1,501 400 ms projections over 180 degrees, in addition to dark-field and white-field scans, using a 2048-by-2048pixel CoolSnap HQ CCD camera (Photometrics, AZ, USA) with a 10X objective lens. Projections were acquired at an energy of 13.8 keV for larval samples to maximize x-ray contrast for PTA-stained samples while compensating for sample thickness (Ding et al., 2019). A 30mm object-scintillator distance was chosen for phase-contrast edge enhancement that most closely resembled contrast visible in histology. Reconstruction was done using the TomoPy toolkit (Gürsoy et al., 2014), including gain and dark-field correction and ring artifact reduction. Reconstruction yielded a 2048 x 2048 x 2048 voxel dataset at an isotropic pixel size of 0.743 µm for larval fish. Whole organism datasets were created by stitching multiple overlapping scans of the same sample. Visualization of data was performed on FIJI for slice-by-slice viewing and simple image transformations such as reslicing across variable angled planes. ITK-Snap was used to manually segment cells for validation sets, providing 3 orthogonal views for confirmation of boundaries as well as a 3D view of segmentations. Avizo was used for all other 3D renderings of datasets.

### 4.4 Blood Cell Identification, Manual Segmentation, and Visual Validation

Blood cells were identified for segmentation by visually comparing micro-CT data to histology (Fig. 3). Even though our manual and automated segmentation pipelines focused on the detection of erythrocytes, the limits of micro-CT imaging resolution for whole zebrafish samples very likely includes lymphocytes and thrombocytes in our detected populations due to their similarity in shape and size. We use the term blood cells as the broadest representation of what is being segmented and characterized. Erythrocytes in zebrafish can be easily identified across either imaging modality, as they are nucleated, and appear as flat, elongated discs. When viewed in cross section, they can have a spear-shaped or spindled appearance. For manual validation sets, regions of interest were chosen based on the apparent variety of blood cell populations and density and included much variation in cell shape and background tissue. In wild-type 5-dpf zebrafish, the heart and posterior dorsal aorta were chosen for these validation regions. Ages of fish were selected to show varying developmental stages and blood cell populations across wild-type and mutant individuals. 4- and 5-dpf wild-type fish were used to show general life stage cell populations. Scanned *hht* mutants at the same 4- and 5-dpf stages were used to compare to the wild-type samples. The rapid cellular degradation of *hht* mutants was expected to manifest a difference in blood cell shape attributes.

4- and 5-dpf wild-type fish are commonly studied developmental ages due to their size and complexity of tissues/cell types (Ronneberger et al., 2012), (Hu et al., 2000). As such, we chose to focus on these stages as a baseline to compare to other wild-type and mutant fish. Additionally, a 5-dpf zebrafish was the first zebrafish to be imaged with ssEM by Hildebrand et al (2017). Blood cells from this 1.7 TB dataset were segmented and used as a comparison for the micro-CT measurements in this study (Fig. 2), ensuring that any morphological metrics derived from the micro-CT data were accurate to their higher-resolution counterpart.

Manual segmentations for automated pipeline validation were performed using the open-source software, ITK-SNAP version 3.8.0 (Yushkevich et al., 2016) (http://www.itksnap.org). Blood cells were segmented in each of three orthogonal views (sagittal, coronal, and axial). Contrast was adjusted in ITK-SNAP to make cell boundaries more visible and distinct from background noise and surrounding tissue. Each cell was segmented slice by slice in ITK-SNAP, and the Crosshair Mode tool was used to view individual cells in each plane. After completion, the segmentation was checked in each plane to evaluate border consistency and limit instances of “bleed-over” inclusion of cells that were near the cell in question. The 3D viewer was used to visualize each cell to check overall shape. The same method was used for the Hildebrand EM cell segmentation. Two such comprehensively segmented manual sets were analyzed for the 5-dpf wild-type sample, to test variability in shape statistics and automatic segmentation accuracy across multiple sections of the zebrafish body.

### 4.5 Automated Blood Cell Segmentation

For automated blood cell segmentation, our approach combines a random forest modeling framework, intensity-based nuclear detection, and a generalized neural net cellular segmentation method to accurately extract blood cells from micro-CT datasets of wild-type and mutant developing fish.

#### 4.5.1 Random Forest Classifier

We utilized the open-source classification software Ilastik to generate a pixel classification model for blood cell detection (Berg et al., 2019). 3D micro-CT training datasets representative of typical scans were used as input data for development of the model. All possible intensity, edge, and texture features were enabled to be used for classification weighing by the algorithm. Training data was provided by manual segmentation of each class, including blood cells, neural nuclei, muscle, pharynx, cartilage, intercellular space, other general soft tissue, and background. Segmentations were added and the model corrected iteratively in real time as Ilastik’s live update feature allows for immediate feedback on changes in the output after any individual segmentation is performed. All data used to train the classifier was completely independent from validation data used to confirm accuracy of the automated segmentation pipeline. Qualitative observation was used to determine when the model was sufficiently developed for generalized blood cell detection from micro-CT images of zebrafish as a preliminary step, and further confirmed by quantitative validation (Fig. 4, Fig. 7, Fig. 8). The model generates a voxel-by-voxel probability map for each processed image, in which the sum of all possible class predictions for every voxel add up to 1. This model, trained on multiple samples, subsequently served as a comprehensive detection method, and was applied to all of the datasets presented in the paper, making use of their similar scanning and reconstruction parameters. This probability map is multiplied by the original data to conserve the intensity range of blood cells while removing background tissue from the image (Fig. 4, Fig. 5).

#### 4.5.2 Cellpose-Based Segmentation

We adapted Cellpose v.1.0.2 (Stringer et al., 2021), a generalized cellular segmentation algorithm, for segmentation of a specific cell type by processing the input data as specified above. To focus the analysis on blood cells in what is otherwise a very background-heavy dataset (including other cell and tissue types not needed for segmentation and potentially interfering with Cellpose), the raw micro-CT data was multiplied by the probability map output from Ilastik before being fed into Cellpose. This retains the values of high probability blood cell volumes close to their original intensities while reducing the signal of other cell and tissue types. The Cellpose output is then further processed by prior biological knowledge through the removal of any segmented objects not fitting with the known shape statistics derived from manually segmented cells taken from validation sets, including volume, flatness, elongation, principal axes length ratios.

Using a Dice-Sorensen measure of overlap to compare to manually segmented sets, it was determined that performing this pre and post processing outperformed Cellpose results from the raw data with no further steps (Fig. 7).

#### 4.5.3 Shape Analysis and Evaluation Metrics

Individual segmented cells were further interrogated by the computation of shape and intensity statistics for more accurate characterization of individual samples and comparisons between developmental stages and genetic conditions. The SimpleITK shape statistic image filter and intensity statistic image filter were used to generate metrics for each cell (Yaniv et al., 2018), (Beare et al., 2018), covering volume, elongation, flatness, equivalent spherical radius, and perpendicular principal 3D axes for shape statistics, as well as average intensity, intensity standard deviation, and intensity skewness for intensity statistics, as described:

**Volume:** Total voxel count per individual cell label.

**Elongation:** A ratio of the moments of every individual cell along the principal long and medium axes.

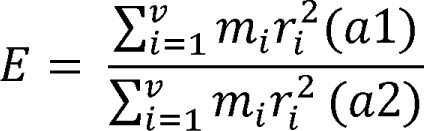

Where v is the total voxel volume of an individual cell, m is a mass constant for every individual voxel, r is the perpendicular distance from a voxel to the axis of rotation, and a1 and a2 are the longest and medium perpendicular axes of the cell, respectively.

**Flatness:** The degree to which each label approximates a mathematical plane. Calculated as a ratio of the moments of every individual cell along the smallest and medium axes.

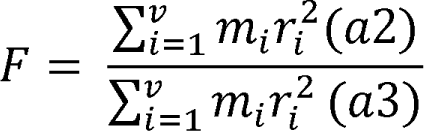

Where v is the total voxel volume of an individual cell, m is a mass constant for every individual voxel, r is the perpendicular distance from a voxel to the axis of rotation, and a2 and a3 are the medium and shortest perpendicular axes of the cell, respectively.

**Equivalent Radius:** Radius of a sphere with the same volume as the cell label.

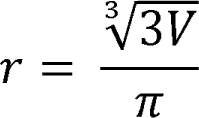

Where V is the total volume of the cell.

**Perpendicular Axes:** Lengths of the three perpendicular axes of each label that encompass the longest and shortest possible distance between points on the cell border.

**Intensity Mean:** The average of the voxel intensities that comprise each label.

**Intensity STD:** The standard deviation of the voxel intensities that comprise each label.

**Intensity Skewness:** The skewness of the voxel intensities that comprise each label.

These values were calculated for every automatically segmented cell and were subsequently compared to the limits of known values derived from manually segmented sets for each genetic and developmental condition, using the full range of obtained variables. All automatically segmented objects outside of these limits were removed as false positives. This ensures higher accuracy in detecting only normal cells across all test populations as high probability areas with unrealistic shapes and intensities are filtered out and additionally ensures consistent application of constraints across all samples.

### 4.6 Segmentation Validation and Experimental Group Comparison

As our analysis places emphasis on automated detection and classification of cells in the context of whole animal 3D datasets, validating the method against both manual sets as well as other methods of automated segmentation is necessary for showcasing utility. Additionally, to characterize blood development across multiple popularly studied stages in development as well as genetic backgrounds, we applied the finalized method to several groups of micro-CT datasets of similarly aged fish. We used the *hht* mutant as a control of abnormal blood development in which we expect heavy differences from the wild-type.

#### 4.6.1 Method Validation

To compare automatically segmented datasets with manually processed data we used the Dice-Sorensen coefficient to calculate overlap between individual cells in the same automatically and manually segmented regions. Setting a threshold for a Dice coefficient between two cells to be considered valid, and treating the manually segmented set as ground truth, we can subdivide all cases of cell overlap between the 2 sets into true positives (TP), false positives (FP), and false negatives (FN). Calculating precision and recall from these values allows for the generation of an F1 score that serves as an indicator of automated segmentation accuracy:

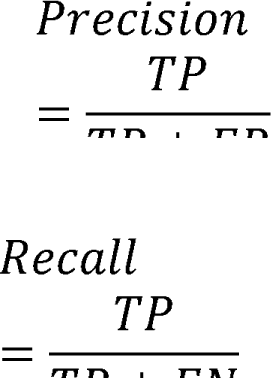

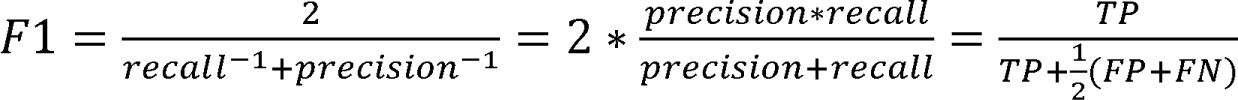

To validate our blood cell detection pipeline against other methods of cellular detection, we compared our automated segmentation results to output obtained from Cellpose, a generalized cell detection model trained on large datasets, run on the raw and unaltered data. All F1 scores were generated by comparing automated methods to manual segmentation, covering three volumes spread across different anatomical regions of the zebrafish we trained our model on, to prevent detection bias that could arise from analyzing only one section with a specific background.

#### 4.6.2 Comparison Methodology

Our preliminary dataset consists of 4 samples, covering 4- and 5-dpf developmental stages as well as wild-type and *hht* genetic backgrounds. Shape analysis of segmentation results yielded tabular data for each sample (4-dpf wild-type, 4-dpf *hht*, 5-dpf wild-type, and 5-dpf *hht*). These data were compiled into csv files with each feature (volume, intensity mean, etc.) as a column and each cell as a row. CSV files were then read into python as data frames using the pandas software package (McKinney, 2010). A label column was added to each cell denoting the genotype and age of each fish from which the cell segmentation mask was extracted. The volume measurement was used for basic data exploration prior to downstream machine learning analysis. Histograms of cell volume were generated with the seaborn package (Waskom, 2021). Blood cell measurement data was then pre-processed for machine learning steps using the standard scaler method in scikit learn. The label and radius columns were removed from the data frames prior to further analysis. The Intensity-associated columns were removed due to image acquisition differences and thus intensity variation between samples. All other measurements were included for linear discriminant analysis (LDA) with 3 components (Fig. 11). LDA was used to visualize and evaluate the ability of the pipeline to resolve different phenotypes.

## Data Availability Statement

Digital histology is publicly available from our Zebrafish Lifespan Atlas (http://bio-atlas.psu.edu) (Cheng, 2004). Raw reconstructed data and images processed through Ilastik, Cellpose, and filtering of the four zebrafish larvae involved in analysis, along with validation data, are available on Dryad (https://datadryad.org/). The used Ilastik model and scripts written for cell detection and characterization are also provided. Due to the large size of these files, image bit depth was scaled down from 32 bit to 8 bit in FIJI prior to uploading. Full bit-depth scans and images, including raw projection data, are available from researchers upon request as a download or transfer to physical media.

The following dataset was generated:

Vanselow, Daniel et al. (2023), Quantitative Geometric Modeling of Blood Cells from X-ray Histotomograms of Whole Zebrafish Larvae, Dryad, Dataset, https://doi.org/10.5061/dryad.f1vhhmh2d The following previously published datasets were used:

Hildebrand, D. G. C., Cicconet, M., Torres, R. M., Choi, W., Quan, T. M., Moon, J., Wetzel, A. W., Scott Champion, A., Graham, B. J., Randlett, O., Plummer, G. S., Portugues, R., Bianco, I. H., Saalfeld, S., Baden, A. D., Lillaney, K., Burns, R., Vogelstein, J. T., Schier, A. F., … Engert, F. (2017). Whole-brain serial-section electron microscopy in larval zebrafish. Nature, 545(7654), 345–349. https://neurodata.io/data/hildebrand17/

## Supporting information

Extra Author Information

## Acknowledgements

The authors thank Dr. Yifu Ding, Dr. Mike Bayerl, Samarth Gupta, Amogh Adishesha, and David Northover for their work, support, and intellectual contributions without which this manuscript would not be possible. Additionally, the authors are very grateful to Dr. Xianghui Xiao and Dr. Francesco De Carlo for their facilitation of synchrotron data acquisition.

