## Supplementary material for "Quantitative Geometric Modeling of Blood Cells from X-ray Histotomograms of Whole Zebrafish Larvae": Extra Author Information

Dear editors,

Thank you so much for considering our manuscript. We attach this supplementary information on the part of the authors to provide clarification on authorship ORCiDs, affiliations, contributions, and a statement on sources for conflicts of interest. We hope that this document will help fill in some of this necessary information and, if necessary, that it will be inserted into the main text wherever it is most appropriate.

Additionally, the Dryad DOI link for our uploaded dataset is still being generated and should be available for use within the next few days. As soon as the manuscript passes review, we will make it publicly available. Please let us know if there is anyone specific that we should send the link to or if we can attach it to these documents post-submission. Finally, we are attaching the four mentioned data tables as separate files and not at the end of the main text as they contain thousands of entries. They are also included in the Dryad dataset. This manuscript was recently submitted to eLife.

Finally, if there is anything else that we can provide to benefit this submission, please reach out to us.

Best regards,

**Maksim A. Yakovlev**

**ORCiD**: 0000-0003-1846-3751

**Affiliations**:

Department of Pathology, Penn State College of Medicine, Hershey, Pennsylvania, USA

The Jake Gittlen Laboratories for Cancer Research, Penn State College of Medicine, Hershey, Pennsylvania, USA

Biomedical Sciences PhD Program, Penn State College of Medicine, Hershey, Pennsylvania, USA

**Contributions:**

- Methodology
- Software
- Validation
- Formal analysis
- Investigation
- Data curation
- Writing 1
- Writing 2
- Visualization

**Ke Liang**

**ORCiD**: N/A

**Affiliations**:

School of Computer, National University of Defense Technology, Changsha, 410073, China

Data Science and Artificial Intelligence Area, College of Information Sciences and Technology, The Pennsylvania State University, University Park, PA 16802, USA

**Contributions:**

- Methodology
- Software
- Validation
- Formal analysis
- Data curation
- Writing 1
- Writing 2
- Visualization

**Carolyn R. Zaino**

**ORCiD**: [0000-0002-4383-6292](https://orcid.org/0000-0002-4383-6292)

**Affiliations**:

Department of Pathology, Penn State College of Medicine, Hershey, Pennsylvania, USA

The Jake Gittlen Laboratories for Cancer Research, Penn State College of Medicine, Hershey, Pennsylvania, USA

**Contributions:**

- Validation
- Investigation
- Resources
- Data curation
- Writing 1
- Writing 2
- Visualization
- Project administration

**Daniel J. Vanselow**

**ORCiD**: [0000-0002-9221-8634](https://orcid.org/0000-0002-9221-8634)

**Affiliations**:

Department of Pathology, Penn State College of Medicine, Hershey, Pennsylvania, USA

The Jake Gittlen Laboratories for Cancer Research, Penn State College of Medicine, Hershey, Pennsylvania, USA

**Contributions:**

- Methodology
- Software
- Validation
- Formal analysis
- Resources
- Data curation
- Writing 2
- Visualization

**Andrew L. Sugarman**

**ORCiD**: [0000-0003-3348-2977](https://orcid.org/0000-0003-3348-2977)

**Affiliations**:

Department of Pathology, Penn State College of Medicine, Hershey, Pennsylvania, USA,

The Jake Gittlen Laboratories for Cancer Research, Penn State College of Medicine, Hershey, Pennsylvania, USA,

Bioinformatics and Genomics PhD Program, Penn State College of Medicine, Hershey, Pennsylvania, USA

**Contributions:**

- Methodology
- Software
- Formal analysis
- Data curation
- Writing 2
- Visualization

**Alex Y. Lin**

**ORCiD**: 0000-0002-1653-4168

**Affiliations**:

Department of Pathology, Penn State College of Medicine, Hershey, Pennsylvania, USA,

The Jake Gittlen Laboratories for Cancer Research, Penn State College of Medicine, Hershey, Pennsylvania, USA

**Contributions:**

- Methodology
- Investigation
- Resources

**Patrick J. La Riviere**

**ORCiD**: [0000-0003-3415-9864](https://orcid.org/0000-0003-3415-9864)

**Affiliations**:

Department of Radiology, The University of Chicago, Chicago, USA

**Contributions:**

- Methodology
- Investigation
- Writing 2
- Supervision

**Yuxi Zheng**

**ORCiD**: N/A

**Affiliations**:

Department of Mathematics, The Pennsylvania State University, University Park, PA, 16802, USA

**Contributions:**

- Methodology
- Supervision

**Justin D. Silverman**

**ORCiD**: 0000-0002-3063-2098

**Affiliations**:

College of Information Sciences and Technology, The Pennsylvania State University, University Park, PA 16802, USA

**Contributions:**

- Methodology
- Writing 2
- Supervision

**John C. Liechty**

**ORCiD**: N/A

**Affiliations**:

Department of Marketing, Pennsylvania State University, State College, PA, 16802 USA

**Contributions:**

- Methodology
- Software
- Supervision

**Sharon X. Huang**

**ORCiD**: [0000-0003-2338-6535](https://orcid.org/0000-0003-2338-6535)

**Affiliations**:

Data Science and Artificial Intelligence Area, College of Information Sciences and Technology, The Pennsylvania State University, University Park, PA 16802, USA

**Contributions:**

- Methodology
- Writing 1
- Writing 2
- Supervision

**Keith C. Cheng**

**ORCiD**: [0000-0002-5350-5825](https://orcid.org/0000-0002-5350-5825)

**Affiliations**:

Department of Pathology, Penn State College of Medicine, Hershey, Pennsylvania, USA

The Jake Gittlen Laboratories for Cancer Research, Penn State College of Medicine, Hershey, Pennsylvania, USA

**Contributions:**

- Conceptualization
- Methodology
- Investigation
- Resources
- Writing 1
- Writing 2
- Supervision
- Project administration
- Funding acquisition

The authors declare that the manuscript and presented data reflect no conflict of interest on any of their parts.
